# Rewiring cattle movements to limit infection spread

**DOI:** 10.1101/2022.08.24.505123

**Authors:** Thibaut Morel-Journel, Pauline Ezanno, Elisabeta Vergu

## Abstract

The cattle tracing databases set up over the past decades in Europe have become major resources for representing demographic processes of livestock and assessing potential risk of infections spreading by trade. The herds registered in these databases are nodes of a network of commercial movements, which can be altered to lower the risk of disease transmission. In this study, we develop an algorithm aimed at reducing the number of infected animals and herds, by rewiring specific movements responsible for trade flows from high- to low-prevalence herds. The algorithm is coupled with a generic computational model describing infection spread within and between herds, based on data extracted from the French cattle movement tracing database (BDNI). This model is used to simulate a wide array of infections, with either a recent outbreak (epidemic) or an outbreak that occurred five years earlier (endemic), on which the performances of the rewiring algorithm are explored. Results highlight the effectiveness of rewiring in containing infections to a limited number of herds for all scenarios, but especially if the outbreak is recent and if the estimation of disease prevalence is frequent. Further analysis reveal that the key parameters of the algorithm affecting infection outcome vary with the infection parameters. Allowing any animal movement from high to low-prevalence herds reduces the effectiveness of the algorithm in epidemic settings, while frequent and fine-grained prevalence assessments improve the impact of the algorithm in endemic settings. According to our results, our approach focusing on a few commercial movements is expected to lead to substantial improvements in the control of a targeted disease, although changes in the network structure should be monitored for potential vulnerabilities to other diseases. Due to its generality, the developed rewiring algorithm could be applied to any network of controlled individual movements liable to spread disease.

## Introduction

Following bovine spongiform encephalopathy and classical swine fever epidemics in the 1990s, the European Union initiated the mandatory identification and registration of cattle in Europe (EU, 2000). This decision led to the creation of national identification databases, such as the cattle tracing system in the United Kingdom (Kao et al., 2006, Vernon, 2011), the French national bovine identification database (BDNI) (Rautureau et al., 2011, Dutta et al., 2014), the Italian national bovine database (Natale et al., 2009, Bajardi et al., 2011) and the database of the Swedish board of agriculture (Nöremark et al., 2009, 2011). These animal tracing systems have enabled the monitoring of infectious livestock diseases and the development of strategies to prevent their spread (Gilbert et al., 2005, Moslonka-Lefebvre et al., 2016, Beaunée et al., 2017), since animal trade is a major transmission pathway between herds. Indeed, commercial exchanges are not only recorded comprehensively, but also controlled by farmers, unlike animal mobility in the wild. These databases, whose reliability has increased over time since their creation (Green and Kao, 2007), are therefore powerful tools for simulating infectious diseases in cattle (Ezanno et al., 2020) and assessing the impact of livestock movements on epidemics (Ezanno et al., 2021).

The information provided by these commercial animal movements can be used as a basis for representing comprehensively the demographic processes and trades between cattle farms located in a given region, using a metapopulation framework (Liu et al., 2007, Widgren et al., 2015). To this end, disease transmission between individuals within a defined set of herds can be modelled, by combining an epidemiological model with existing data on births, deaths and movements. This type of models accounts at least for two ways of spreading the infection: by contact within a herd, or by actually moving animals between herds. This is for instance the case for paratuberculosis, a cattle disease mainly spread between herds by trade (Beaunée et al., 2015, Biemans et al., 2021). Manipulating the structure of cattle movement is expected to have a direct impact on the latter and an indirect impact on the former.

The structure of these trade movements can be understood through the prism of graph theory: herds are the vertices of a commercial exchange network, whose edges are the movements of livestock (Dubé et al., 2009). Thus, each herd can be characterised using graph metrics, e.g. its in- and out-degree, i.e. the number of herds it has respectively bought animals from and sold animals to. Network-based control strategies then aim to modify the structure of the network to reduce infection risks. Removing vertices (Rautureau et al., 2011, Büttner et al., 2013) or edges (Yang et al., 2013, Green et al., 2009) through trade ban or culling is a method used to slow down epidemics. In a context of cattle exchange however, preventing farmers from buying or selling livestock entails high economic costs. Therefore, this strategy cannot be used routinely or over extended periods of time. It is likely better suited to the management of regulated diseases, the consequences of which are also very costly and for controlling outbreaks of newly introduced diseases. Conversely, the application of such drastic methods on the longer term for endemic diseases may not be feasible.

Edge rewiring is a less radical approach able to balance the trade-off between health risks and economic costs. This method corresponds to the modification of one or both vertices that an edge connects (Gross et al., 2006, Piankoranee and Limkumnerd, 2020, Britton et al., 2016, Ball and Britton, 2020). Although most of the theoretical literature on the subject rather considers rewiring in the context of human contact networks, it has also been used to study epidemic spread in cattle movement networks (Gates and Woolhouse, 2015, Mohr et al., 2018, Ezanno et al., 2021, Biemans et al., 2022). For instance, Gates and Woolhouse (2015) present a rewiring method that creates an entirely new movement network disconnecting large buyers from large sellers, while retaining the total number of animals bought or sold by each herd. This method requires information at the network level, the criteria used being the distributions of in- and out-degrees of all herds. Global-level information is also generally required for most rewiring methods in contact networks, although Piankoranee and Limkumnerd (2020) proposed a method based on local information. In their study, rewiring is decided at the vertex level, according to its status and those of its direct neighbours. Controlling cattle movements depending on the sanitary status of their origin has been proposed in previous studies, e.g. by Hidano et al. (2016). Their study presents different scenarios regarding farmers’ practices, especially their tendency to avoid buying cattle from regions with a higher incidence of bovine tuberculosis. The approach presented here is similar, albeit at a finer grain: preventing farmers from buying cattle from herds with a higher prevalence of the target disease.

This study presents a new rewiring method to reduce the spread of infections in a cattle movement network. To do this, we developed a rewiring algorithm aimed at preventing the movements of animals from higher-prevalence herds to lower-prevalence ones. It was based on an edge-level criterion: the estimated difference in prevalence between the herd of origin and the herd of destination of the movement considered. For this study, we tested the algorithm in conjunction with a computational epidemiological model describing the spread of a nonspecific disease, whose infectiousness was parametrically defined. The impact of the algorithm was tested using a real commercial movement network, based on dataset from the French cattle tracing system (BDNI). In contrast with similar rewiring approaches developed recently to target specific diseases (Ezanno et al., 2021, Biemans et al., 2022), we propose a more generalist approach aimed at investigating the effectiveness of this type of method in a broader context. After presenting the movement network used as an example, the model and the algorithm, we consider various outputs of simulations with and without rewiring, concerning the functioning of the algorithm itself, its impact on infection propagation, and on the structure of the cattle movement network.

## Data and methods

### Cattle movement network

In order to test the algorithm on a actual network of commercial bovine movements, we use an extraction from the French national bovine identification database (BDNI). It includes all cattle herds in Brittany (a French region) that sold or bought at least one animal during the year 2014. This set of 21,548 herds is referred to as the ‘metapopulation’ thereafter. Every animal in the dataset is included regardless of breed or age, in order to have a larger number of movements per herd over this period of time. Three types of commercial exchanges are considered: (i) ‘internal movements’ have an origin and a destination among the herds in the dataset, (ii) ‘imports’ have only a destination in the dataset and (iii) ‘exports’ have only an origin in the dataset. They represent respectively 64%, 16% and 20% of the commercial exchanges involving at least one herd of the dataset. Each commercial exchange of animals is assumed to take place directly from one herd to another, neglecting intermediaries. This means that markets and sorting centres are not considered for this study. They differ from herds in that they tend to concentrate a large number of animals, but for a limited period of time (less than a day for markets, a few days for sorting centres). In addition, the dataset also includes information about the demographic events in the herd, which are considered as a special type of movements: (iv) births have only a destination, corresponding to the herd where the animal is born, and (v) deaths have only an origin, corresponding to the last herd recorded for the animal.

The dataset is represented as a network with herds and internal movements corresponding to the vertices and edges, respectively. This network is (i) dynamic, i.e. movements are characterised by the date at which they occur, (ii) weighted, i.e. a single edge represents the set of all movements from herd *A* to herd *B*, with a weight corresponding to the number of movements, and (iii) directed, i.e. movements from herd *A* to herd *B* are accounted for separately from movements from herd *B* to herd *A*. The network therefore includes 21,548 vertices and 100,088 edges. The total number of internal movements over 2014 is 206,640, thus the average edge weight is 2.06.

### Epidemiological model: within and between-herd dynamics and infection settings

The model developed aims to simulate pathogen transmission within herds, and infection spread between herds through cattle movements. A full description of the model is available in Supplementary material 1. The model is stochastic in discrete time – each time-step corresponding to a day of 2014 – and in discrete space – by integrating the network of herds and movements described above. Commercial exchanges and demography are data-based: movement *m* is characterised by its origin *O*_*m*_, its destination *D*_*m*_, its date according to the dataset 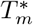 and the date at which it is simulated *T*_*m*_. By default, movements are simulated according to the dataset, i.e. 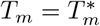. Within-herd dynamics are based on a SIRS model with three parameters: the infection rate *β*, the recovery rate *γ* – therefore the average infection duration is 1*/γ* – and the rate of return to susceptibility *δ*. At each time-step *t*, herd *h* is characterised by its number of susceptible, infected and recovered individuals, noted respectively *S*_*h*_(*t*), *I*_*h*_(*t*) and *R*_*h*_(*t*). The total herd size *N*_*h*_(*t*) is defined as the sum of these three values and infection prevalence as *P*_*h*_(*t*) = *I*_*h*_(*t*)*/N*_*h*_(*t*).

Each simulated infection begins with an initial outbreak in a metapopulation without infection, i.e. with only susceptible individuals. At *t* = *t*_*I*_, the date of the outbreak, 10% of all herds in the metapopulation are infected, by replacing 1 susceptible individual with 1 infected individual in each of the herds. The probability of a herd being part of this 10% is proportional to the number of imports in the herd according to the 2014 dataset. The rationale is that herds receiving the most individuals from herds outside of the metapopulation are the most likely to introduce a new infection.

Two types of infections are considered for the study: epidemic and endemic. An infection is defined as ‘epidemic’ if it starts at the outbreak, i.e. if *t*_0_ = *t*_*I*_. The initial state of the infection is then as described above. An infection is defined as ‘endemic’ if its start date is five years after the outbreak, i.e. *t*_0_ = *t*_*I*_ +1825 days. The initial state of infection is then the result of a five-year infection, simulated using the same epidemiological model and an extraction from the BDNI over Brittany between 01/01/2009 and 31/12/2013. Endemic simulations for which the infection goes extinct before *t*_0_ are discarded, so that only initial states that are not disease-free are considered.

### Prevalence status of the herds

The algorithm developed aims at identifying and preventing movements of cattle ‘at risk’, i.e. those from higher-prevalence herds to lower-prevalence herds. The differences in prevalence are based on prevalence classes, numbered from 1 to *c*. Class *i* corresponds to prevalence values between *b*_*i*_ and *b*_*i*+1_, with the lowest boundary *b*_1_ = 0 and the highest boundary *b*_*c*+1_ = 1. The prevalence status of herd *h* at time *t*, noted 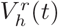, is then the class including its prevalence, i.e. 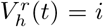 if *P*_*h*_(*t*) ∈ [*b*_*i*_; *b*_*i*+1_], and 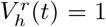 if *P*_*h*_(*t*) = 1. Yet, this ‘real’ prevalence status is not the one used by the algorithm. Rather, it uses an ‘observed’ prevalence status, noted 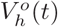, which is recorded at *t*_*obs*_ and then remains the same for *q* time-steps., i.e. 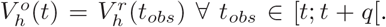. No additional error on the observed status (e.g. because of imperfect test specificity or sensitivity) is assumed, so that it always corresponds to the real prevalence status at *t*_*obs*_. Movements are considered ‘at risk’ if the observed prevalence status of their origin is strictly greater than that of their destination.

### Sequential rewiring

The algorithm works by permuting the origins of pairs of movements, one of which is at risk, so that neither of them is at risk after the rewiring. The pairs of movements are created such that 1 ≤ *c*_*ON*_ ≤ *c*_*DR*_ *< c*_*OR*_ ≤ *c*_*DN*_ ≤ *c*, with *c*_*OR*_ and *c*_*DR*_ the observed status of the origin and destination of the movement at risk and *c*_*ON*_, *c*_*DN*_ those of the origin and destination of the other movement. By permuting the origins, the algorithm creates a movement with an origin of status *c*_*ON*_ and a destination of status *c*_*DR*_, and another movement with an origin of status *c*_*OR*_ and a destination of status *c*_*DN*_. Then neither of the two movements is at risk, since *c*_*ON*_ ≤ *c*_*DR*_ and *c*_*OR*_ ≤ *c*_*DN*_.

For all movements to occur at a given time-step, the algorithm performs these permutations in a specific order to ensure that no potential rewiring is missed. Supplementary material 2 describes this functioning of the algorithm over a single time-step in pseudo-code. Firstly, it defines all possible quadruplets of prevalence classes *{c*_*OR*_, *c*_*DR*_, *c*_*ON*_, *c*_*DN*_ *}*. These quadruplets are arranged primarily in ascending order of *c*_*DR*_, secondarily in descending order of *c*_*OR*_, thirdly in ascending order of *c*_*ON*_ and fourthly in descending order of *c*_*DN*_. This order ensures that no potential permutation is missed by the algorithm. For each quadruplet, the algorithm then permutes the origins of *k* pairs of movements, with *k* the minimum between the number of movements at risk and the number of other movements considered.

Once all possible permutations are performed, there might be remaining movements at risk set to be performed on this time-step. Firstly, these movements are postponed to the next day, to be potentially rewired with another set of movements. The postponed movements are then prioritised for rewiring on the following day. Yet, postponing commercial movement represents a constrain for farmers. Therefore, a maximal delay during which a movement can be postponed Δ_*MAX*_ is fixed for the algorithm. Thus, remaining movement *m* is postponed to the next day only if it was not already postponed Δ_*MAX*_ days, i.e. if 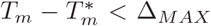. If the algorithm prohibits any movement at risk, the remaining movements that cannot be postponed (called ‘problematic’ movements) are replaced by one export with the origin of the problematic movement as origin and one import with the destination of the problematic movement as destination. Otherwise, the problematic movement is conserved as such. Overall, the algorithm therefore depends on four parameters: the number of prevalence classes *c*, the period at which observed status is updated *q*, the maximum delay Δ_*MAX*_ and whether movements at risk are prohibited.

### Simulations

Simulations are performed on the dataset between 01/01/2014 (defined as *t* = 0) and 01/01/2015 (*t* = 365). Different epidemiological settings are explored by manipulating the SIRS model parameters (*β, γ* and *δ*) and infection type (epidemic or endemic). Two clustering analyses are performed on the preliminary simulations to define six epidemiological settings (Supplementary material 3): weak, moderate and strong epidemic settings and weak, moderate and strong endemic settings (Fig. S2).

The effectiveness of the algorithm is tested by running simulations with 3 × 3 × 3 × 2 combinations of the algorithm parameters, respectively (i) the number of prevalence classes *c* (2, 3 or 4 classes), (ii) the update period *q* (1, 28 or 91 days), the maximum delay Δ_*MAX*_ (1, 3 or 7 days) and (iv) the prohibition of movements at risk (yes or no). Each combination, as well as a control without rewiring, are simulated 100 times for each of the six epidemiological settings.

Preliminary simulations are also carried out for each epidemiological setting between 01/01/2009 (*t* = −1825) and 31/12/2013 (*t* = −1), with an initial outbreak at *t*_*I*_ = −1825. On the one hand, the number of susceptible, infected and recovered individuals of each herd at *t* = −1 are used as the starting numbers for the endemic simulations (starting at *t* = 0). On the other hand, the boundaries of the prevalence classes *b*_*i*_ used by the algorithm are set as quantiles of the distribution of prevalence values. These boundaries ensure that the number of herds of each class is roughly the same at the start of the simulation. If fewer than 1*/c* herds have a null prevalence, *b*_*i*_ is the ((*i* − 1) */c*)^*th*^ quantile of the distribution. If it is greater than 1*/c, b*_1_ = *b*_2_ = 0 and *b*_*i*_ is the ((*i* − 2) */* (*c* − 1))^*th*^ quantile of the distribution.

### Outcomes and analyses of numerical explorations

The simulations outcomes are listed in Table 1. They are related either to (i) the functioning of the algorithm, (ii) the infection or (iii) the network of internal movements modified by the algorithm.

**Table 1:** List of the outcomes computed from the simulations. The infection-related outcomes were computed for each simulation separately. The algorithm and network-related ones were computed for each simulation with the algorithm.

| Outcomes related to | Notation | Description |
| --- | --- | --- |
| Algorithm | $n_{rew}(t)$ | Number of movements rewired at time $t$ |
| | $n_{del}(t)$ | Number of delayed movements at time $t$ |
| | $n_{prob}(t)$ | Number of problematic movements at time $t$ |
| | $n_{risk}(t)$ | Number of movements at risk at time $t$ |
| | $n_{err}(t)$ | Number of movements undetected as at risk at time $t$ |
| Infection | $n_{inf}$ | Number of herd infections |
| | $n_{ext}$ | Number of herds in which the infection goes extinct |
| | $a_{dur}$ | Average duration of infection |
| | $c_{inc}(t)$ | Cumulative incidence at time $t$ |
| | $n_{herd}(t)$ | Number of infected herds at time $t$ |
| | $n_{ind}(t)$ | Number of infected individuals in the metapopulation at time $t$ |
| | $a_{prev}(t)$ | Average prevalence in the infected herds at time $t$ |
| Network | $n_{SCC}$ | Number of strongly connected components |
| | $max_{SCC}$ | Size of the largest strongly connected component |
| | $ind_h$ | In-degree of herd $h$ |
| | $outd_h$ | Out-degree of herd $h$ |

The algorithm-related outcomes *n*_*rew*_(*t*), *n*_*del*_(*t*) and *n*_*prob*_(*t*) are computed each time-step after rewiring, while *n*_*risk*_(*t*) and *n*_*err*_(*t*) are computed before. These latter outcomes are computed by using the real prevalence status of the herds, rather than the observed ones. A movement *m* is included in *n*_*risk*_(*t*) if 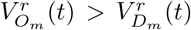, and also included in *n*_*err*_(*t*) if 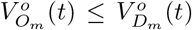 at the same time. The proportion of undetected movements at risk is computed on a weekly basis, to account for intra-week variability in the number of livestock movements. Over week *w*, this proportion *p*_*err*_(*w*) is:

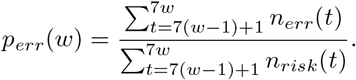

The Spearman’s correlation coefficient *ρ* between *p*_*err*_(*w*) and the number of weeks since last update (from 1 to 4 weeks if *q* = 28 days, from 1 to 13 weeks if *q* = 91 days) is also computed to assess the relationship between errors in herd prevalence status and time. The Spearman’s coefficient is preferred because it does not assume any particular distribution of the involved variables.

The impact of the algorithm on the infection dynamic is estimated through *c*_*inc*_(*t*), i.e. the cumulative number of herds newly infected over the simulation. The variations in *n*_*herd*_(*t*) and *n*_*ind*_(*t*) over time are also presented in Supplementary material 4. Besides, the overall impact of the algorithm on the infection is assessed using a global multivariate sensitivity analysis, following Lamboni et al. (2011) and using the *multisensi* package of the *R* software (Bidot et al., 2018), which is used to perform sensitivity analyses on a multivariate output. For this analysis, twelve variables are derived from the infection-related outcomes. The three outcomes computed once per simulation *n*_*inf*_, *n*_*ext*_ and *a*_*dur*_ are used as such. In addition, the maximum, minimum and final values over the whole period simulated (respectively noted *max*(*u*(*t*)), *min*(*u*(*t*)) and *u*(365) for outcome *u*(*t*)) of *n*_*herd*_(*t*), *n*_*ind*_(*t*) and *a*_*prev*_(*t*) are also computed. The analysis includes a principal component analysis (PCA) on the scaled variables, which are used as the multivariate output for the sensitivity analysis. Two generalised sensitivity indices (GSI), which are weighted means of the sensitivity indices over all the dimensions of the PCA, are computed for each algorithm parameter: the total index (tGSI) including interactions with other parameters, and the first-order index (mGSI), not including them. The first principal component of the PCA is also used to assess the distribution of the simulations depending on the algorithm parameters.

The network-related outcomes are based on an static view of the network aggregating all the internal movements performed during the simulation, from *t* = 0 to *t* = 365. Therefore, they take into account the rewiring performed by the algorithm, and the potential removal of problematic movements if movements at risks are completely prohibited. The outcomes recorded for the modified networks are compared to the same metrics for the original network defined by the 2014 dataset. The strongly connected components – from which *n*_*SCC*_ and *max*_*SCC*_ are computed – correspond to groups of vertices linked to each other by a directed path. The percentiles of the distributions of *ind*_*h*_ and *outd*_*h*_ of all herds in the static network are used to assess the in-degree and out-degree distributions, respectively.

## Results

### Outcomes related to the algorithm

Our results show that number of movements rewired varies greatly depending on the date of the outbreak. It is negligible in the epidemic settings, with 80% of simulations with a total of rewired movements between 192 (fewer than 0.1% of all movements) and 2250 (1.1%). However, it is larger in the endemic settings, with 80% of simulations with between 17,344 (8.4% of all movements) and 33,640 (16.3%) movements rewired. Besides, increasing the value of Δ_*MAX*_ logically increases the number of delayed movements (which is 0 by definition for Δ_*MAX*_ = 0) and decreases the number of problematic movements. In the endemic settings, the problematic movements represent a small proportion of the movements detected as high risk (median: 5.4%, 9^*th*^ decile: 17.4%). In the epidemic settings however, they represent a larger part (median: 14.3%, 9^*th*^ decile: 59.7%), although their absolute numbers remain low (median: 129, 9^*th*^ decile: 651). Because of the overwhelming number of initially non-infected herds in these simulations, the movements at risk are likely more difficult to rewire, and thus more likely to be tagged as problematic by the algorithm.

Increasing the herd status update period *q* is not associated with a decrease in the number of rewiring events (Fig. 1A, 1B). The value of *q* is even rather positively correlated with the number of rewiring events in epidemic settings. This suggests that the algorithm performs more erroneous rewiring as *q* increases. This is confirmed by the distributions of Spearman’s correlation coefficient between *p*_*err*_(*w*) and the number of weeks since last update *ρ* with *q* = 91 days (Fig. 1D), in epidemic settings (80% of values of *ρ* between -0.01 and 0.50) and in endemic settings (80% of values of *ρ* between 0.39 and 0.75). This is also somewhat the case with *q* = 28 days (Fig. 1C), although the correlations are weaker, in endemic (80% of values of values between -0.09 and 0.79) as well as in epidemic settings (80% of values of values between -0.05 and 0.34).

**Figure 1:**
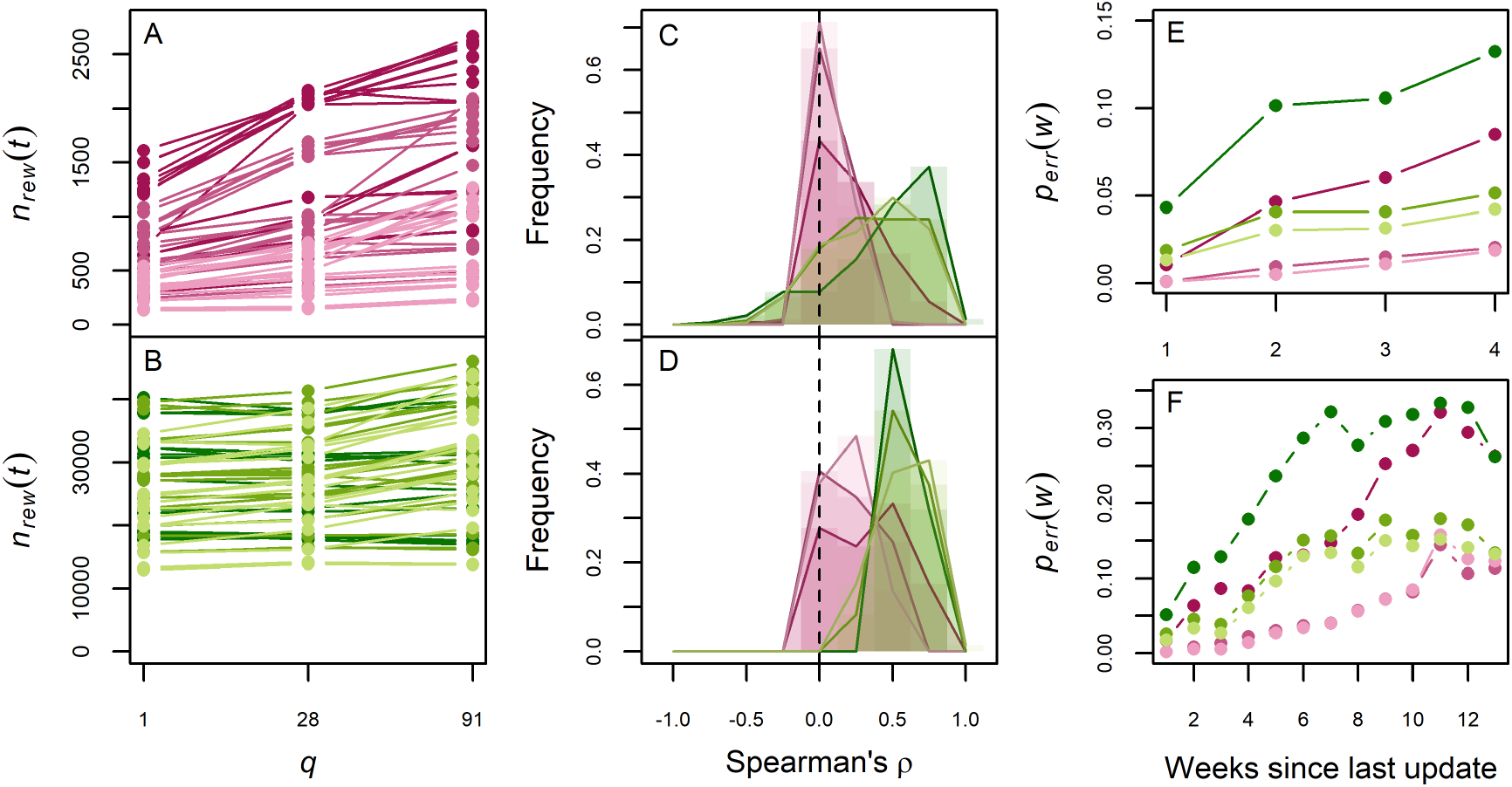
Impact of the update period *q* on the undetected movements at risk, in epidemic (magenta) or endemic settings (green), weak (light), moderate (medium) or strong (dark). First column: total number of rewiring events as a function of the update frequency *q*, averaged over all simulations for a same algorithm parameter combination, in epidemic (A) and endemic settings (B). Second column: distribution of Spearman’s correlation coefficients (*ρ*), with *q* = 28 days (C) and *q* = 91 days (D). Third column: average proportion of undetected movements at risk *p*_*err*_(*w*) as a function of the number of weeks since the last update, with *q* = 28 days (E) and *q* = 91 days (F).

The average proportions of undetected movements at risk *p*_*err*_(*w*) all tend to increase with the number of weeks since the last update *w* (Fig. 1E, 1F). This increase is systematically greater for the largest value of *q*, up to *p*_*err*_(*w*) = 0.3. However, they also appear to have reach a plateau after 10 weeks. This suggests that a further increase in the update period *q* would not strongly increase the proportion of undetected movements at risk. As for Spearman’s correlation coefficient *ρ*, the increase is greater in endemic settings than in epidemic settings.

### Outcomes related to the infection

Comparison of the results with and without rewiring shows the overall effectiveness of the algorithm in containing the infection (Fig. 2). Regardless of the epidemiological setting and the combination of parameters considered, the cumulative number of herds newly infected *c*_*inc*_(*t*) remains systematically lower after rewiring. The algorithm is particularly effective in weak and moderate epidemic settings, where very few herds are infected during the year. In other epidemiological settings, the impact of the algorithm varies more strongly depending on the scenario considered. Results for *n*_*herd*_(*t*) and *n*_*ind*_(*t*) are presented in Supplementary material 4. In epidemic settings, variations in *n*_*herd*_(*t*) logically follow closely those of *c*_*inc*_(*t*). Hence, the algorithm also reduces the increase in the total number of infected herds. It also reduces the total number of infected individuals, although the impact is not as strong as for herds. In endemic settings, the value of *n*_*herd*_(*t*) remains similar during the whole simulation without rewiring (Fig. S3), despite new infections according to variations in *c*_*inc*_(*t*). This indicates a turnover in the infection at the metapopulation level, with populations losing the infection through the acquisition of resistance or the culling and trade of infected animals. By reducing the number of new infections, the algorithm actually therefore reduces the total number of infected herds over time. However, its impact is smaller on the total number of infected individuals (Fig. S4).

**Figure 2:**
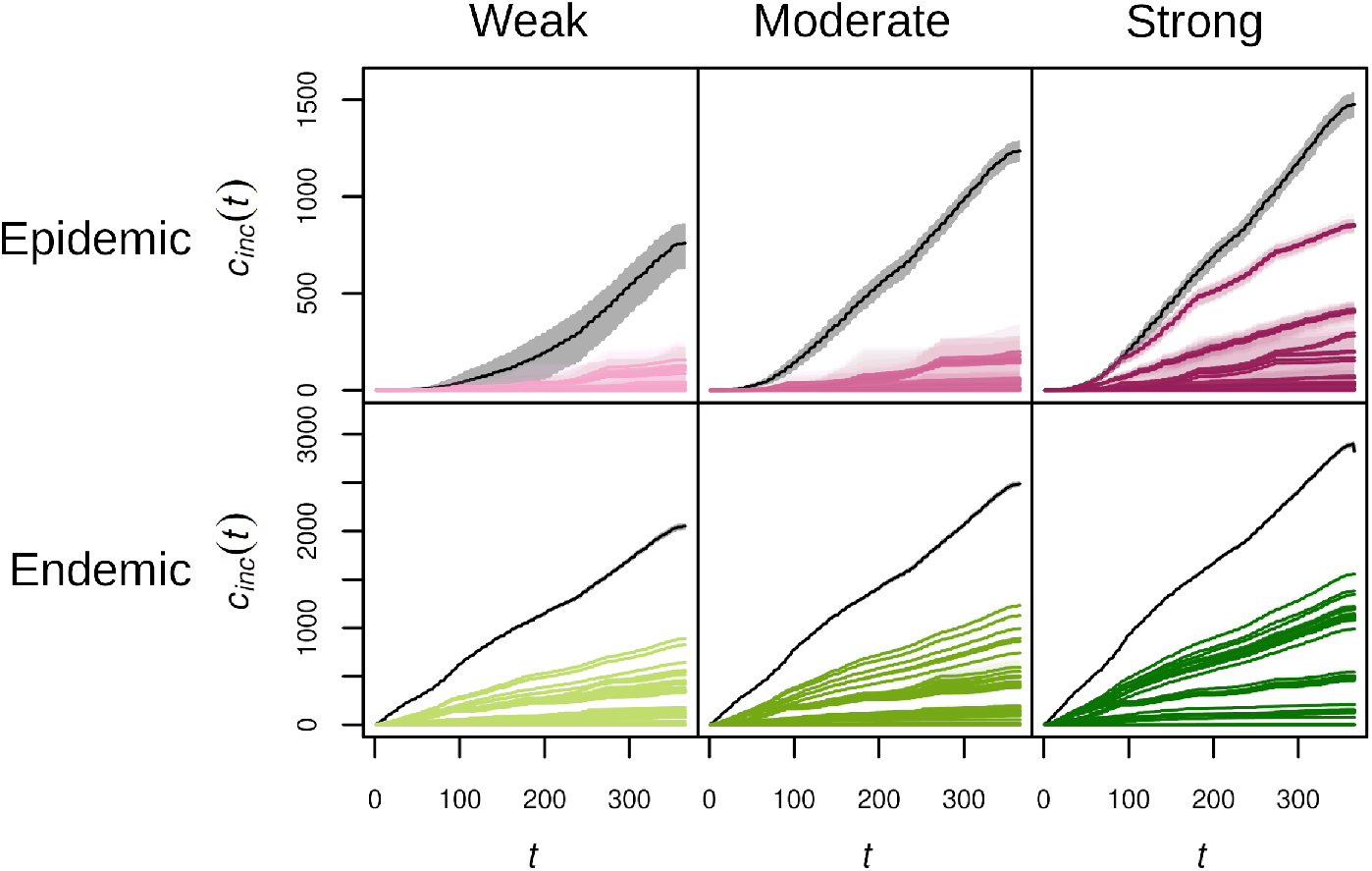
Cumulative incidence *c*_*inc*_(*t*), in number of herd infections, as a function of time (*t*, in days), for simulations with (colour) or without rewiring (black), in epidemic (1^*st*^ row, magenta) or endemic settings (2^*nd*^ row, green), weak (1^*st*^ column, light), moderate (2^*nd*^ column, medium) and strong (3^*nd*^ column, dark). Each combination of algorithm parameters is represented by its mean over the repetitions (solid line) and an interval of 80% of simulations (envelope).

The sensitivity analysis shows differences in the relative importance of the algorithm parameters on the reduction of the infection (Fig. 3). Three different patterns of sensitivity to the algorithm parameters are observed. Firstly, simulations in weak and moderate epidemic settings exhibit an overwhelming sensitivity to the prohibition of movements at risk. Secondly, those in strong epidemic or endemic settings exhibit a strong sensitivity to the number of prevalence classes *c*. Finally, those in weak and moderate endemic settings exhibit a more balanced sensitivity to all parameters, with a substantial difference between total and first-order indices for the maximum delay Δ_*MAX*_, the number of classes and the prohibition of movements at risk. These differences suggest an interaction between the three algorithm parameters. Besides, simulations for every epidemiological setting are somewhat sensitive to the update period *q*.

**Figure 3:**
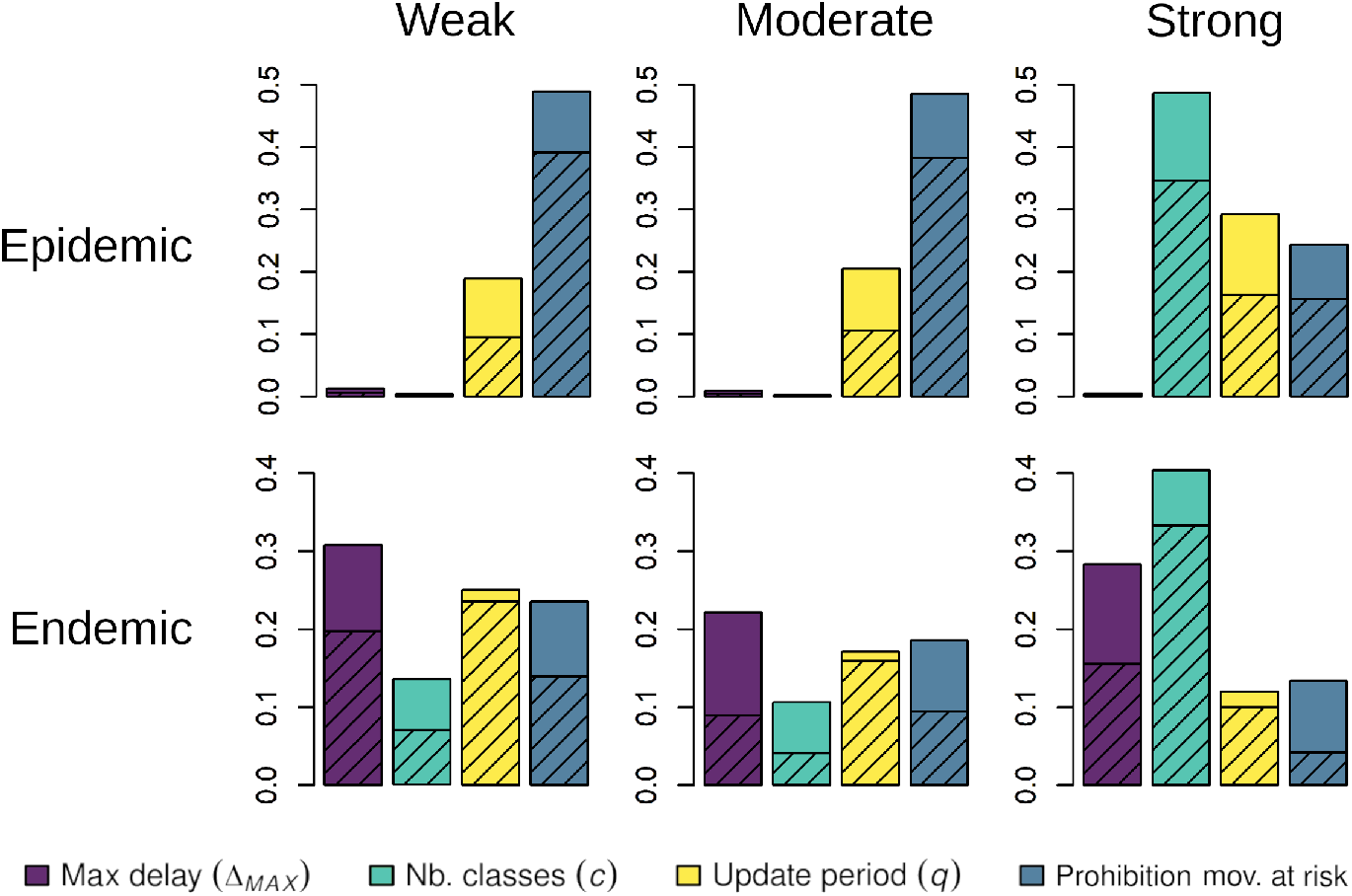
Generalised sensitivity indices (GSI) of the maximum delay Δ_*MAX*_ (purple) the number of prevalence classes *c* (cyan), the update period *q* (yellow) and the prohibition of movements at risk (blue), in epidemic (1^*st*^ row) or endemic settings (2^*nd*^ row), weak (1^*st*^ column), moderate (2^*nd*^ column) and strong (3^*rd*^ column). The total indices (tGSI) are in solid colour and the first-order indices (mGSI) are hatched.

The PCA performed as a first step of the sensitivity analysis is used to explore further the way algorithm parameters impact the infection-related outputs. Supplementary material 5 shows that the first principal component of the PCA is globally positively correlated with outputs describing the extent of the infection. The distributions of simulations along this first principal component therefore provides information about the way algorithm parameter values affects the extent of the infection. Supplementary material 6 presents these distributions for every epidemiological setting and every algorithm parameter, while Fig. 4 displays some of the most relevant distributions. Fig. 4A shows that, in the weak epidemic setting, simulations in which movements at risk are prohibited almost always score lower on the first principal component than those in which they are not. The distribution is similar in the moderate epidemic setting (Fig. S6), which has similar sensitivity indices (Fig. 3). Interestingly, distributions of simulations in strong epidemic or endemic settings show that those with *c* = 2 score higher on their respective first component, while those with *c* = 3 and *c* = 4 are not different (Fig. 4B, 4F). A similar pattern is observed with the maximum delay in the weak endemic setting: only simulations with Δ_*MAX*_ = 0 score higher on the first principal component (Fig. 4D). In the strong epidemic setting, the two high-scoring peaks in the distribution according to *c* (Fig. 4B) correspond to the simulations with *q* = 28 and *q* = 91 (Fig. 4C), highlighting an interplay between the number of classes *c* and the update period *q*. No interplay between Δ_*MAX*_ and *q* is visible in the weak endemic setting, although Fig. 4E show that the score of simulations on the first principal component is positively correlated with *q*. Distributions in the moderate endemic setting are similar to those in the weak endemic setting (Fig. S6).

**Figure 4:**
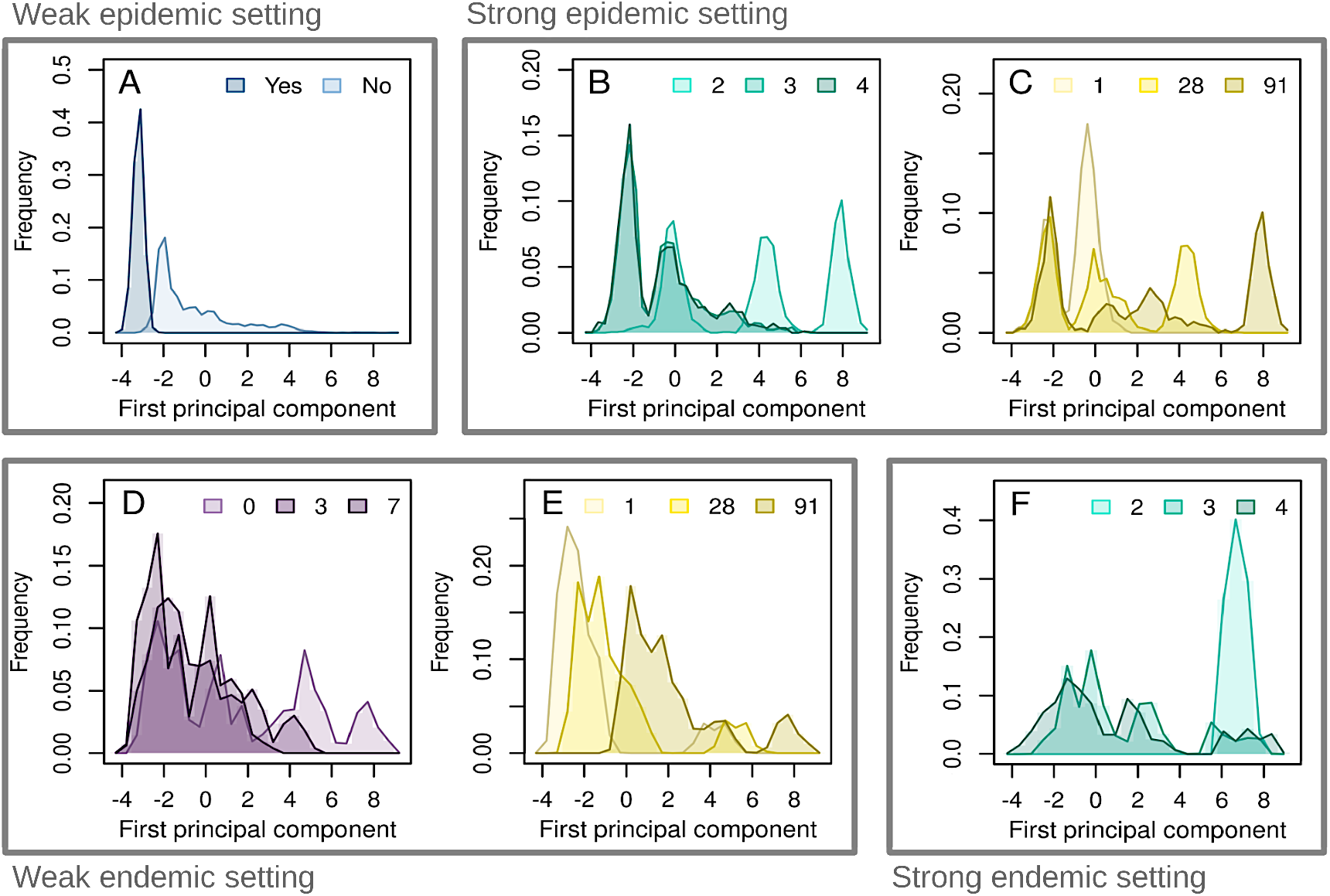
Distribution of the simulations on the first component of the PCA performed as a first step of the sensitivity analysis, in the weak epidemic setting (A), the strong epidemic setting (B, C), the weak endemic setting (D, E) and the strong endemic setting (F). The outputs are divided by maximum delay (purple, D), management of problematic movements (blue, A), number of prevalence classes (cyan, B and F) and herd status update period (yellow, C and E).

### Outcomes related to the movement network

In endemic settings, rewiring movements increase the in- and out-degrees of the herds, i.e. the number of different herds they are connected to (see Supplementary material 7). The increase is small but systematic, for every algorithm parameter value (Fig. S7). In addition, the algorithm also affects the strongly connected components of the network in endemic settings. On the one hand, the algorithm reduces their number, all the more that the infection was strong (Fig. 5). On the other hand, the size of the largest strongly connected component is increased in most, but not all simulations (64%, 67% and 80% of simulations in low, moderate and high endemic settings, respectively). It should be noted that the lesser impact of the algorithm on the network in epidemic settings can be explained by a number of rewiring events 25 times smaller on average than in endemic settings.

**Figure 5:**
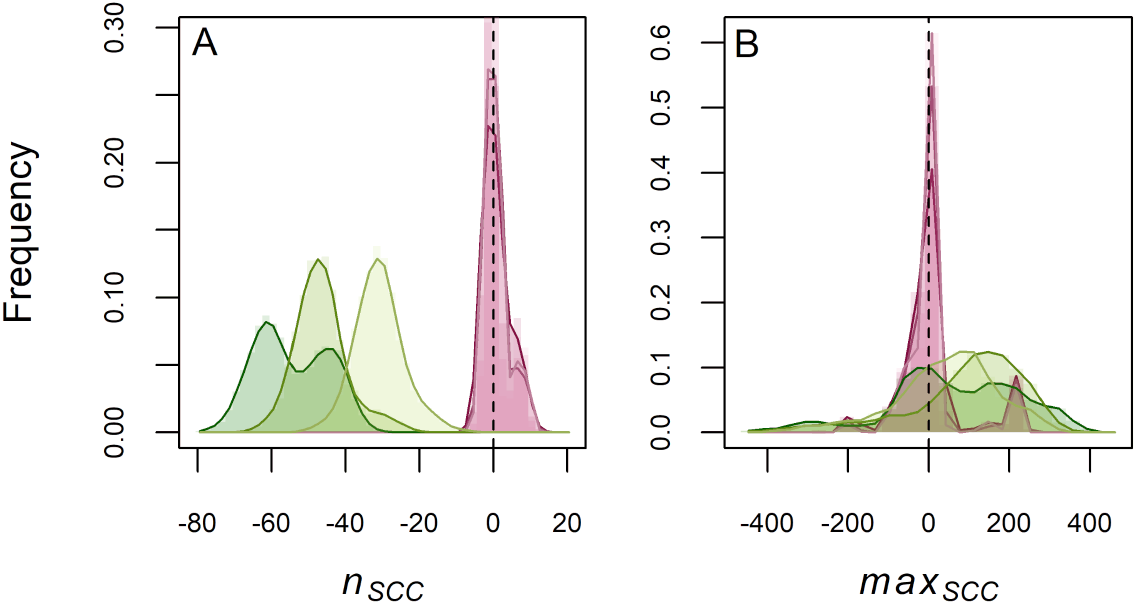
Distributions of the differences in number of strongly connected components (A, *n*_*SCC*_) and in size of the largest strongly connected component B, *max*_*SCC*_) between rewired networks and the original one from the dataset, for epidemic (magenta) and endemic (green) settings, weak(light), moderate (medium) and strong (dark).

## Discussion

The rewiring algorithm we developed for this study is able to reduce the extent of infections, in the absence of any other restriction measure and for a large panel of disease parameters (infection rate *β*, recovery rate *γ* or rate of return to susceptibility *δ*). However, the extent of the reduction varies between the different epidemiological settings considered. Indeed, infections are almost completely prevented with weak or moderate epidemic settings, while they still develop or persist for other settings, although not as much as without any rewiring. However, the decrease in the number of infected herds is not necessarily coupled with a decrease in the number of infected individuals. This result highlights the tendency of the algorithm to concentrate infected individuals in the already infected herds. The algorithm therefore performs a trade-off that is beneficial to the metapopulation as a whole – with fewer infected herds – but detrimental to the smaller number of already infected herds, in such a situation where movement rewiring is not combined with complementary on-farm measures to reduce within-herd infection prevalence. This is the case for the infections in an epidemic setting, in which the prevalence in the infected herds increases over the year. This is also the case for infections in endemic settings, in which new sensitive individuals could still be born or imported.

The sensitivity analysis on the infection-related outcomes reveals that the impact of the parameters of the algorithm is highly dependent on the epidemiological setting. Prohibiting the movements at risk, i.e., removing the movements that cannot be rewired and are delayed as much as possible, is mostly significant if the infection is not too strong and is just beginning. Only in these cases can the infection be fully contained by the prohibition. Increasing the maximal delay improves the performance of the algorithm in an endemic setting, for which the number of movements rewired is much larger than in epidemic settings. In those, delaying the movement to the next day increases substantially the opportunities for rewiring. The other two parameters are both related to the definition of the prevalence statuses used by the algorithm. A greater number of prevalence classes, which mainly impacts rewiring during strong infections, improves the separation of disease-free herds from the rest. Indeed, considering more prevalence classes lowers the upper boundary of the lowest one, which included only herds with very few or no infected animals, thus allowing the algorithm to effectively protecting disease-free herds. A longer update period between updates of the prevalence status makes the rewiring algorithm more error-prone, with a proportion of undetected movements at risk increasing with the time since the last update, at least up to ten weeks. This result is visible for any epidemiological setting, suggesting that any increase in the frequency of update to the status of the herds should improve the effectiveness of the algorithm. Conversely, the results indicate that increasing the number of prevalence classes to more than two, or having a maximum delay greater than zero, improves the efficiency of the algorithm much more than further increases.

As expected, the impact of rewiring on the commercial movements network structure is limited, as it targeted a few movements only: less than 20% of the movements for endemic infections and less than 2% of them for epidemic infections. Nevertheless, rewiring tends to increase the overall connectedness of the herds during endemic infections. Indeed, the increase in degree and in size of the largest strong component indicates that the algorithm has connected herds that were originally not so. These metrics are generally correlated with higher expected epidemic risks (Kiss et al., 2006, Dubé et al., 2009). The use of such a rewiring method to manage actual bovine movements should take into account this potential increase in the risk of spreading other diseases. The algorithm could be extended to assess multiple diseases at once, but the additional constraints on rewiring would likely reduce its effectiveness.

The main hurdle to implementing this rewiring method in a real-life setting is its reliance on accurate and frequent prevalence data from a large number of farms. Firstly, a lack of specificity or sensitivity in the tests used might lead to an overestimation or underestimation of the prevalence in the herds, depending on the disease considered. Although we show that the algorithm remains efficient even though the observed prevalence differed from the real ones, this additional error could add up with the one observed in our study. However, the impact of such errors is also expected to be mitigated by the use of prevalence classes, so that small differences do not necessarily change the prevalence status of the herds. Secondly, obtaining frequent prevalence data for a large number of farms remains challenging. Bulk milk-based sampling systems could be used for some diseases in cattle (e.g. Garoussi et al., 2008, Humphry et al., 2012, with bovine viral diarrhoea), which would facilitate prevalence estimation for multiple herds at once, thus reducing the associated costs. It would also be possible to reduce the sampling effort by focusing on a subset of herds to monitor. Firstly, this sampling effort should take into account additional information available thanks to measures already in place. For instance, the status of some herds could be approximated through health accreditation schemes (e.g. Ezanno et al., 2021), with herds already identified as disease-free could be automatically assigned to the lowest prevalence status for a given duration. Secondly, herds to monitor could be selected based on their role in disease spread, notably through network metrics. Indeed, central herds in the movement network, i.e. those through which a large proportion of animal movements pass, are expected to play a larger role in the spread of infection (Rautureau et al., 2011, Natale et al., 2011). Hoscheit et al. (2021) reviewed centrality measures taking into account the dynamic nature of the movement network, based on the BDNI. They found that the TempoRank index would for example be a good candidate for selecting a subset of herds to be specifically monitored and taken into account by the algorithm.

In this study, we use a network corresponding to commercial movements between every farm in Brittany (an administrative region of France) over a year to test the efficiency of the algorithm. The choice to limit the size of the network is notably motivated by computational limitations. Indeed, simulating a stochastic spread of the disease on a national scale over six years - five for the preliminary simulations and one for the main simulations - would have been considerably more costly, thus limiting the exploration of variations in the parameters of the SIRS model and the algorithm. Yet, this choice had additional implications that should be underlined.

Firstly, a substantial proportion of the movements involve herds outside of Brittany and are therefore not concerned by the rewiring. Indeed, 20% of all movements whose destination was in the metapopulation had an origin outside of it. In our simulations, these imports are assumed to not be movement at risk, i.e. that the prevalence status of their origin is never higher than that of their destination. This is not trivial, as it presumes that imports do not create greater infection risks than internal movements. In a real-life context, applying this rewiring method in a single region would therefore require an additional management of the risk associated with imports. Yet, extending its use nationally should mitigate this problem, as the proportion of imports is expected to be much lower at this scale. Secondly, every commercial movement between farms is considered to test the algorithm, regardless of breed or age, in order to have a large enough set of movements. Indeed, additional criteria, concerning for instance the breed of the animals, could be added easily by providing the algorithm with movements for individuals in each category separately. However, such criterion would reduce the rewiring possibilities of the algorithm and therefore its effectiveness. Again, the network of commercial movements at the national scale could be large enough to separate the movements by breed or consider only movements of specific breeds.

Although the algorithm is tested on historical data from the BDNI for this study, it could also be used prospectively as part of decision-making tools, barring the limitations presented above. Indeed, the rewiring method could work without any simulation of infection, if herd statuses were provided otherwise. Given these statuses and the potential movements to occur, the algorithm would also suggest necessary changes to prevent movements at risk. In this context, the implementation of these changes would also depend on the actual decision of the informed farmers. Unless rewiring is enforced, it is expected that constraints other than sanitary ones would affect movements, which would impact the effectiveness of the algorithm. Coupling it with a decision-making model could provide additional insight on this impact. In order to make it easier to use as part of such decision-making tools, the algorithm has been specifically designed to be able to include additional, different constraints.

Besides, the rewiring method presented is not limited to cattle, but applicable to a much wider range of networks in animal and plant populations, e.g. among seed exchange networks, which face similar infection risks (Jeger et al., 2007, Pautasso et al., 2010). While the need for controlled movements makes this method more relevant to agricultural systems, the spatial and temporal scales considered can also be adapted depending on the context. Indeed, the daily time-step and the region level were used here as they correspond to the BDNI data structure, but are not necessary for the algorithm to work. The usefulness of our rewiring method could therefore extend beyond cattle concerns, even though the effectiveness of the algorithm in other contexts remains to be tested.

This study demonstrates the effectiveness of a rewiring method targeting specific movements to reduce infection risks. Our approach thus differs radically from that presented by Gates and Woolhouse (2015), as it also aims at generating minimal changes in the structure of the movement network. However, this study builds upon the results from Ezanno et al. (2021), by confirming the effectiveness of this method beyond the specific case of bovine paratuberculosis. Indeed, the algorithm presented by Ezanno et al. (2021) and later by Biemans et al. (2022), was developed specifically to address the control of bovine paratuberculosis, notably characterised by an endemic status and a low detection rate. To do so, they used a specific age-structured epidemiological model (Camanes et al., 2018) and an algorithm calibrated to target the disease. This was also the case for instance of Mohr et al. (2018), which specifically targeted foot-and-mouth disease. Conversely, the present study aims at assessing more comprehensively the effectiveness of the algorithm. It is tested for different epidemiological settings – both endemic and epidemic – using a non-specific epidemiological model, and for broad range of parameter values. This study is therefore complementary to the previous ones, by bringing a broader perspective on the impact of rewiring in animal movement network on infectious diseases in general.

## Abbreviation

BDNI: Base de données nationale d’identification animale

## Data, scripts, code, and supplementary information availability

The code for the algorithm, as well as additional scripts for formatting the data or running preliminary simulations and dummy test data, are freely available at https://sourcesup.renater.fr/projects/pub-rewir-algo/. The dataset used in the study is an extraction from the French national bovine identification database (BDNI), which is confidential, and therefore cannot be provided publicly.

