## Supplementary material for "Rewiring cattle movements to limit infection spread"

This supplementary material provides additional information about the article: *Rewiring cattle movements to limit infection spread* (Authors: T. Morel-Journel, P. Ezanno, E. Vergu). It includes the following:

1. Description of the metapopulation epidemiological simulation model
2. Description of the rewiring algorithm in pseudo-code
3. Definition of the epidemiological settings
4. Variations in the number of infected herds and individuals during the simulations
5. PCA as a first step of the multivariate sensitivity analysis
6. Distributions of the simulations on the first PCA axes depending on the algorithm parameters
7. Changes in in- and out-degree distributions in the movement network after rewiring

### 1. Description of the metapopulation epidemiological simulation model

The simulation model used in the article describes the variations in the number of susceptible, infected and recovered individuals in each herd in the metapopulation at each date. It is stochastic and discrete in time and space. Each time-step (equal to a day) comprise a movement phase including the simulation of commercial exchanges and demographic events, and a transition phase corresponding to the simulation of infections, recoveries or returns to susceptibility.

*The movement phase*

As stated in the main text of the article, the term 'movement' is used here to refer to commercial exchanges as well as births and deaths of individuals. The five types of movements are always simulated in the following order: births, imports, internal movements, exports and deaths. This order has been chosen so that individuals are only added to the metapopulation before the movements and only removed afterwards. As there is no information about the actual order of movements within the same day, this ensures that there are enough individuals in the herds to perform the internal movements. Within each set of movements of the same type, the order used follows the list of movements provided by the database.

Each birth and import is simulated by adding an individual to the destination, each export and death by removing an individual from the origin. Internal movements are simulated by removing an individual from the origin and adding an individual to the destination.

The status of each moved individual is randomly drawn from a multinomial distribution with $n = 1$ and probabilities of being susceptible $p_S(m)$, being infected $p_I(m)$, and being recovered $p_R(m)$:

$$(M_S, M_I, M_R) \sim Multinomial(1, p_S(m), p_I(m), p_R(m)). \tag{1}$$

Therefore, the status of the individual moved is susceptible if $M_S = 1$, infected if $M_I = 1$ and recovered if $M_R = 1$. One individual is moved at a time, in order to avoid drawing more individuals of a given status than available in the origin.

The newborns are always susceptible, meaning that $p_S(m) = 1$, while $p_I(m) = p_R(m) = 0$ if $m$ is a birth. For internal movements, exports and deaths, the origin of the movement $O_m$ is a herd of the metapopulation, in which every animal has an equal chance of being chosen. Therefore, the probabilities for an animal of exiting herd $O_m$ at time $T_m$ are defined as follows:

$$\begin{array}{l} p_S(m) = S_{O_m}(T_m)/N_{O_m}(T_m) \\ p_I(m) = I_{O_m}(T_m)/N_{O_m}(T_m) \\ p_R(m) = R_{O_m}(T_m)/N_{O_m}(T_m) \end{array} . \tag{2}$$

For imports, the proportions of susceptible, infected and recovered individuals of the origin are unknown. Instead, a phantom herd is created by pooling the herds of the metapopulation together, and used as an origin as described above. If there is no rewiring during the simulation, every herd is included into this phantom herd. If there is a rewiring event during the simulation, herd $h$ is included in the phantom herd created for an import $m$ only if its observed prevalence status is lower than or equal to the one of the destination of the movement, i.e. if $V_h^o(T_m) \leq V_h^{D_m}(T_m)$.

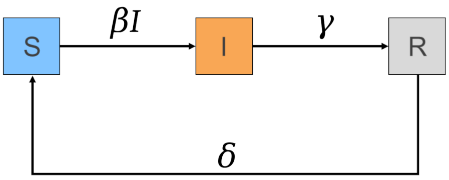


**Figure S1**: Functioning of the SIRS model, with susceptible(S), infected(I) and recovered (R) compartments, and the transition rates between the three states.

*The transition phase*

The intra-herd phase is defined by a SIRS model with three parameters: the infection rate $\beta$, the recovery rate $\gamma$ and the rate of return to susceptibility $\delta$ (Figure S1). The number of newly susceptible ($S_h^{'}(t)$), infected ($I_h^{'}(t)$) and recovered individuals ($R_h^{'}(t)$) in herd $h$ is drawn from binomial distributions:

$$\begin{array}{l} S_h^{'}(t) \sim Binomial(R_h^m(t), \delta) \\ I_h^{'}(t) \sim Binomial(S_h^m(t), \beta I_h^m(t)) \\ R_h^{'}(t) \sim Binomial(I_h^m(t), \gamma) \end{array} , \qquad (3)$$

with $S_h^m(t)$, $I_h^m(t)$ and $R_h^m(t)$ the numbers of susceptible, infected and recovered individuals in herd $h$ after the simulation of movements, respectively. The number of susceptible, infected and recovered individuals at $t+1$ is then computed as follows:

$$\begin{array}{l} S_h(t+1) = S_h^m(t) + S_h^{'}(t) - I_h^{'}(t) \\ I_h(t+1) = I_h^m(t) + I_h^{'}(t) - R_h^{'}(t) \\ R_h(t+1) = R_h^m(t) + R_h^{'}(t) - S_h^{'}(t) \end{array} . \qquad (4)$$

#### 2. Description of the rewiring algorithm in pseudo-code

A given simulated time-step $t$ is separated into three successive steps: (1) the potential update of the observed herd statuses, (2) the rewiring of internal movements and (3) the simulation of the epidemiological model. This section presents the functioning of step 2 in pseudo-code.

Inputs fixed for the simulation

Given:
  a number of estimated prevalence classes $c$ (integer $> 0$)
  a maximal delay $\Delta_{MAX}$ (integer $\geq 0$)
  a boolean indicating whether movements at risk were prohibited **PROHIB** (true or false)

Permutation of the origins

For $c_{DR}$ from $1$ to $(c-1)$, in ascending order:
  For $c_{OR}$ from $c$ to $(c_{DR}+1)$, in descending order
    For $c_{ON}$ from $1$ to $c_{DR}$, in ascending order:
      For $c_{DN}$ from $c$ to $c_{OR}$, in descending order:
        Set **lRisk** the list of all movements $mR$ such that $V^r_{O_{mR}}(t) = c_{OR}$ and $V^r_{D_{mR}}(t) = c_{DR}$
        Set **lNorm** the list of all movements $mN$ such that $V^r_{O_{mN}}(t) = c_{ON}$ and $V^r_{D_{mN}}(t) = c_{DN}$
        Set **minLen** the smallest value between the lengths of **lRisk** and of **lNorm**:
        If **minLen** $> 0$:
          For $k$ between 1 and **minLen**:
            Set **mRisk** the $k^{th}$ movement in **lRisk**
            Set **mNorm** the $k^{th}$ movement in **lNorm**
            Set **NewOR** the origin of **mNorm**
            Set **NewON** the origin of **mRisk**
            Change the origin of **mRisk** to **NewOR**
            Change the origin of **mNorm** to **NewON**

Management of remaining movements at risk

Set **lRemain** the list of movements $mE$ such that $V^r_{O_{mE}}(t) > V^r_{D_{mE}}(t)$
Set **lToDelay** the list of movements in **lRemain** $mD$ such that $T_{mD} < (T^*_{mD} + \Delta_{MAX})$
Set **lProblem** the list of movements in **lRemain** $mP$ such that $T_{mP} = (T^*_{mP} + \Delta_{MAX})$
For **mD** in **lToDelay**:
  Increase $T_{\mathbf{mD}}$ by $1$

If **PROHIB**
  For movement **mP** in **lProblem**:
    Set **newImport** as an import with $D_{\mathbf{newImport}} = D_{\mathbf{mP}}$ and $T_{\mathbf{newImport}} = t$
    Set **newExport** as an export with $O_{\mathbf{newExport}} = O_{\mathbf{mP}}$ and $T_{\mathbf{newExport}} = t$
    Replace **mP** by **newImport** and **newExport**

### 3. Definition of the epidemiological settings

The six epidemiological settings presented in the main text of the article are defined according to two clustering analyses. These analyses are performed on sets of simulated infections in the metapopulation presented in the main text ('Cattle movement network' in 'Data and methods'), comprising all cattle herds in Brittany between 01/01/2014 and 31/12/2014. The first clustering analysis is performed with an epidemic infection type, i.e. with an outbreak starting at the beginning of the simulation $t_I = t_0$. The second is performed with an endemic infection type, i.e. with an outbreak starting five years prior to the simulation $t_I = t_0 - 1825$ days. Three clusters are identified for each set of simulations, using the k-means method. Each set of simulations includes $5 \times 5 \times 5$ combinations of values of $\beta$ ($1.10^{-4}$, $2.5.10^{-4}$, $5.10^{-4}$, $7.5.10^{-4}$, $1.10^{-3}$), $\gamma$ and $\delta$ ($1.10^{-3}$, $2.5.10^{-3}$, $5.10^{-3}$, $7.5.10^{-3}$, $1.10^{-2}$ for each). The 125 combinations of parameter values are simulated 500 times each.

The infection-related outcomes considered are the one described in Table 1 of the main text of the article. Twelve variables are derived from these outcomes. The three outcomes computed once per simulation $n_{inf}$, $n_{ext}$ and $a_{dur}$ are considered as such. In addition, the maximum, minimum and final values of $n_{herd}(t)$, $n_{ind}(t)$ and $a_{prev}(t)$ are computed. They are respectively noted $max(u(t))$, $min(u(t))$ and $u(365)$ for outcome $u(t)$.

The 12 variables used for the clustering analysis are computed for each run of each setting and scaled, i.e. centred and divided by their standard deviation over all simulations. For both infection types (endemic and epidemic), the three clusters correspond to three levels of infection intensity: weak, moderate and strong (Figure S2). However, the parameters driving the structuring of the clusters are different between the two infection types. For the epidemic infection type, clusters are mainly structured according to the $\beta$ value (positively correlated with the strength of the infection). The strong epidemic setting also differs from the others in having an overall lower value of $\gamma$ (Table S1). For the endemic infection type, clusters are mainly structured according to the $\gamma$ value (negatively correlated with the strength of the infection). In addition, the

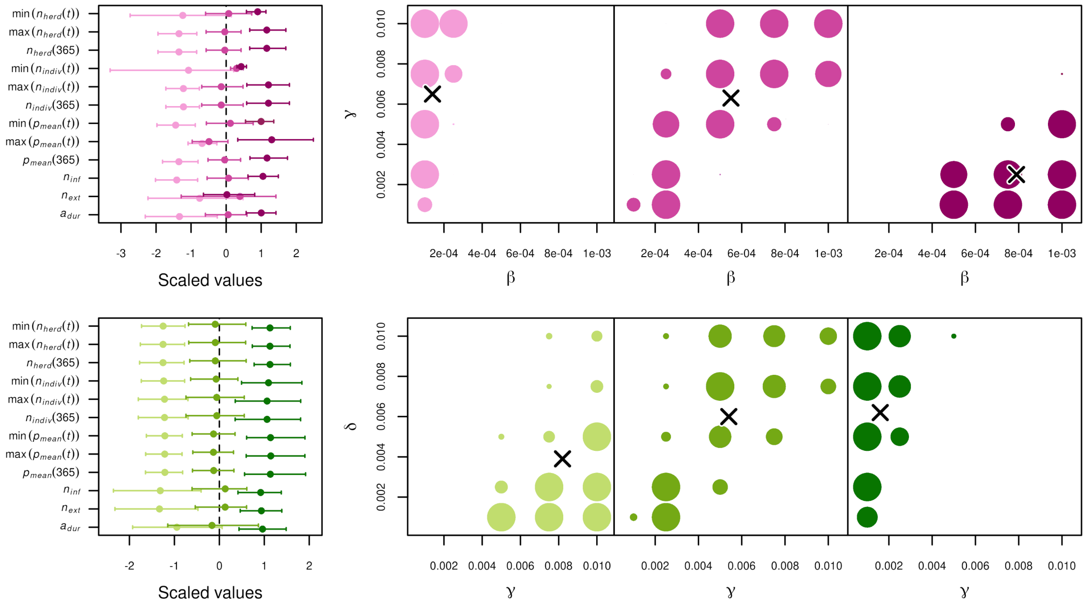


**Figure S2**: Clustering analyses for simulations with an endemic ($1^{st}$ row , magenta) or an epidemic infection type ($2^{nd}$ row, green). Left: Mean and 80% of the scaled variables for the simulations in the cluster of weak (light), moderate (medium) and strong (dark) setting. Right: distribution of the simulations in each cluster according to the values of $\beta$ and $\gamma$ (for epidemic settings) or the values of $\gamma$ and $\delta$ (for endemic settings). The size of the dots corresponds to the proportion of simulations in the settings. The crosses correspond to the values of the parameters averaged over all the simulations belonging to the cluster.

| Epidemiological setting | | $\beta(\times 10^{-3})$ | $\gamma(\times 10^{-3})$ | $\delta(\times 10^{-3})$ |
|---|---|---|---|---|
| Epidemic | Weak | 0.14 | 6.5 | 5.1 |
| | Moderate | 0.55 | 6.3 | 5.5 |
| | Strong | 0.79 | 2.5 | 5.5 |
| Endemic | Weak | 0.43 | 8.2 | 3.9 |
| | Moderate | 0.55 | 5.4 | 6.0 |
| | Strong | 0.58 | 1.6 | 6.2 |

**Table S1**: Average values of $\beta$, $\gamma$ and $\delta$ for the simulations belonging to each cluster identified.

weak endemic setting has overall lower $\delta$ values.

#### 4. Variations in the number of infected herds and infected individuals in the simulations

The results presented here show the variation in the number of infected herds $n_{herd}(t)$ and infected individuals $n_{ind}(t)$ during the simulations, with and without rewiring, for each epidemiological setting. Simulations with rewiring are performed for scenario (i.e. combination of algorithm parameters) presented in the main text

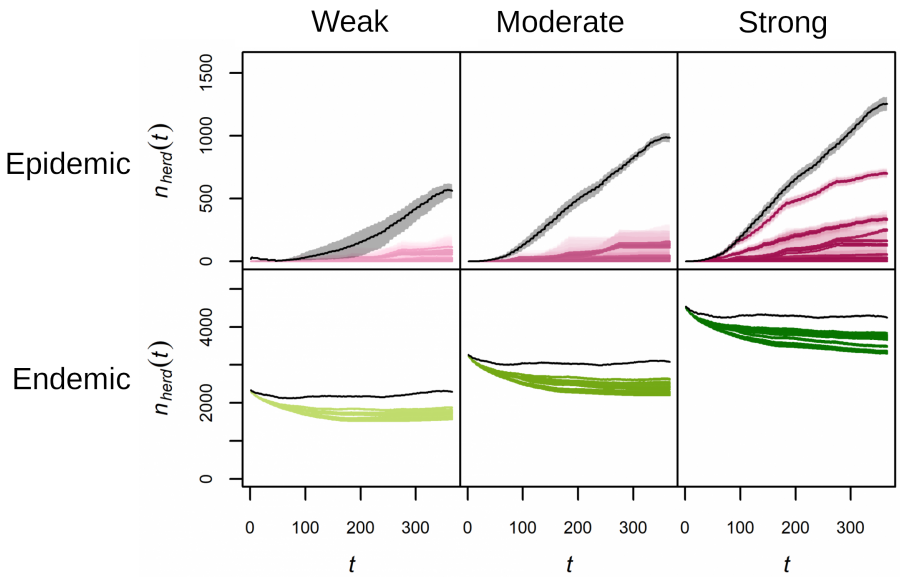


**Figure S3**: Number of infected herds $n_{herd}(t)$ as a function of time ($t$, in days), for simulations with (colour) or without rewiring (black), in epidemic ($1^{st}$ row, magenta) or endemic ($2^{nd}$ row, green) settings, weak (light), moderate (medium) or strong (dark). Each scenario (algorithm parameter combination) is represented by its mean over the repetitions (solid line) and and an interval of 80% of simulations (envelope)

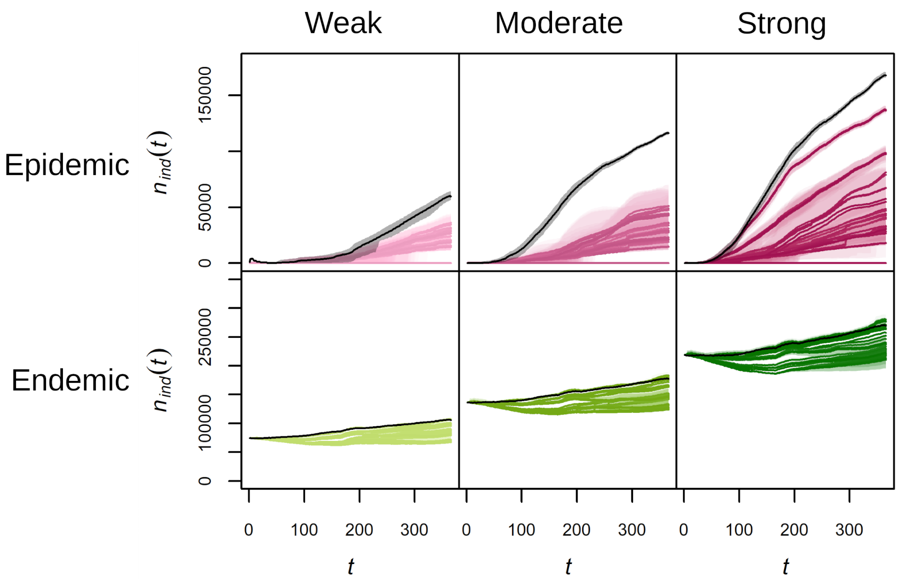


**Figure S4**: Number of infected individuals $n_{ind}(t)$ as a function of time ($t$, in days), for simulations with (colour) or without rewiring (black), in epidemic ($1^{st}$ row, magenta) or endemic ($2^{nd}$ row, green) settings, weak (light), moderate (medium) or strong (dark). Each scenario (algorithm parameter combination) is represented by its mean over the repetitions (solid line) and and an interval of 80% of simulations (envelope)

of the article. The values of $n_{herd}(t)$ (Figure S3) show a substantial impact of the algorithm in reducing infection, regardless of the scenario. The algorithm slows down the increase in number of infected herds, even reaching $n_{herd}(t) = 0$ for some scenarios in epidemic settings. Besides, the algorithm also reduces $n_{herd}(t)$ in endemic settings, while it remains broadly constant during simulations without rewiring.

The values of $n_{ind}(t)$ (Figure S4) show a similar, albeit smaller, improvement brought about by the rewiring. The number of infected individuals is also lower for all scenarios in epidemic settings, although the difference is not as pronounced. However, this is not always the case in endemic settings, meaning that, for some scenarios, the algorithm only concentrates infected cattle into a smaller number of herds, without reducing the total number of infected animals.

### 5. PCA as a first step of the multivariate sensitivity analysis

The multivariate sensitivity analysis following Lamboni et al. (2011) includes a preliminary principal component analysis (PCA) on the outputs, which are described in the main text of the article. While the analysis takes all the components of the PCA into account, the first principal component (FPC) is specifically investigated in the following. Its inertia is presented in Table S2 for each epidemiological scenario, and shows that it systematically captures a majority – but not all – of the variability in the data analysed.

| Epidemiological setting | | Inertia of the $1^{st}$ component of the PCA |
|---|---|---|
| Epidemic | Weak | 55.56% |
| | Moderate | 63.87% |
| | Strong | 70.73% |
| Endemic | Weak | 68.58% |
| | Moderate | 64.08% |
| | Strong | 68.82% |

**Table S2**: Inertia of the first principal component of the PCA prior to the sensitivity analyses.

The correlation of the FPC with each output is very similar for each epidemiological setting (Fig. S5). The correlations observed suggest that this FPC describes well the extent of the simulated infection. Indeed, it is clearly positively correlated with outcomes corresponding to more extensive infections ($n_{herd}$, $n_{indiv}$, $n_{inf}$). Exceptions include $max(n_{herd}(t))$ in endemic settings and $min(n_{herd}(t))$ and $min(n_{herd}(t))$ in epidemic settings, which are never correlated with the FPC. This is expected, as the number of infected herds always decreases with rewiring in endemic settings (see Fig. S3), and the number of infected herds and individuals always increase in epidemic settings (see Fig. S3 and Fig. S4). These values therefore correspond to those at $t = 0$, which are the same (or close to) across all simulations. The correlation between the FPC and the outcomes concerning prevalence ($n_{prev}$) is not as clear, but also overall positive.

However, the FPC is positively correlated to $n_{ext}$ and negatively correlated to $a_{dur}$, which is unexpected. This can be explained by their strong link with $n_{inf}$, the total number of infections of herds (Table S3). First, $n_{ext}$ is always strongly positively correlated with $n_{inf}$, as the extinction of the infection in a herd can

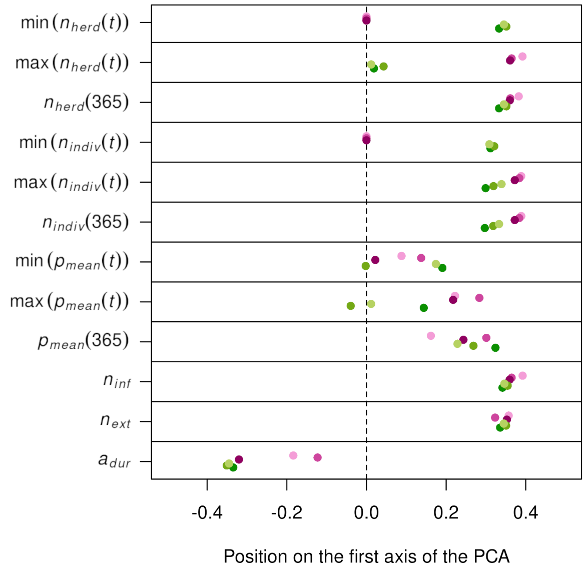


**Figure S5**: Position of the twelve infection-related outputs on the first principal component of the PCA prior to the sensitivity analysis, for the weak (light), moderate (medium) and strong (dark) epidemic (magenta) and endemic (green) settings.

only occur after the infection of this herd. Secondly, $a_{dur}$ is negatively correlated with $n_{inf}$, meaning that short-lived infections are more likely if many herds are infected. This could be the result of the algorithm: by concentrating infected individuals in fewer herds, the algorithm creates few, long-lived infections, compared to the infections observed when the algorithm was less effective.

| Epidemiological setting | | $cor(n_{inf},\ n_{ext})$ | $cor(n_{inf},\ a_{dur})$ |
|---|---|---|---|
| Epidemic | Weak | 0.85 | -0.44 |
| | Moderate | 0.83 | -0.32 |
| | Strong | 0.98 | -0.78 |
| Endemic | Weak | 0.99 | -0.99 |
| | Moderate | 0.98 | -0.98 |
| | Strong | 0.98 | -0.97 |

**Table S3**: Pearson's correlations between $n_{inf}$ and $n_{ext}$or $a_{dur}$ (scaled values) for each epidemiological scenario.

#### 6. Distributions of the simulations on the first PCA axes depending on the algorithm parameters

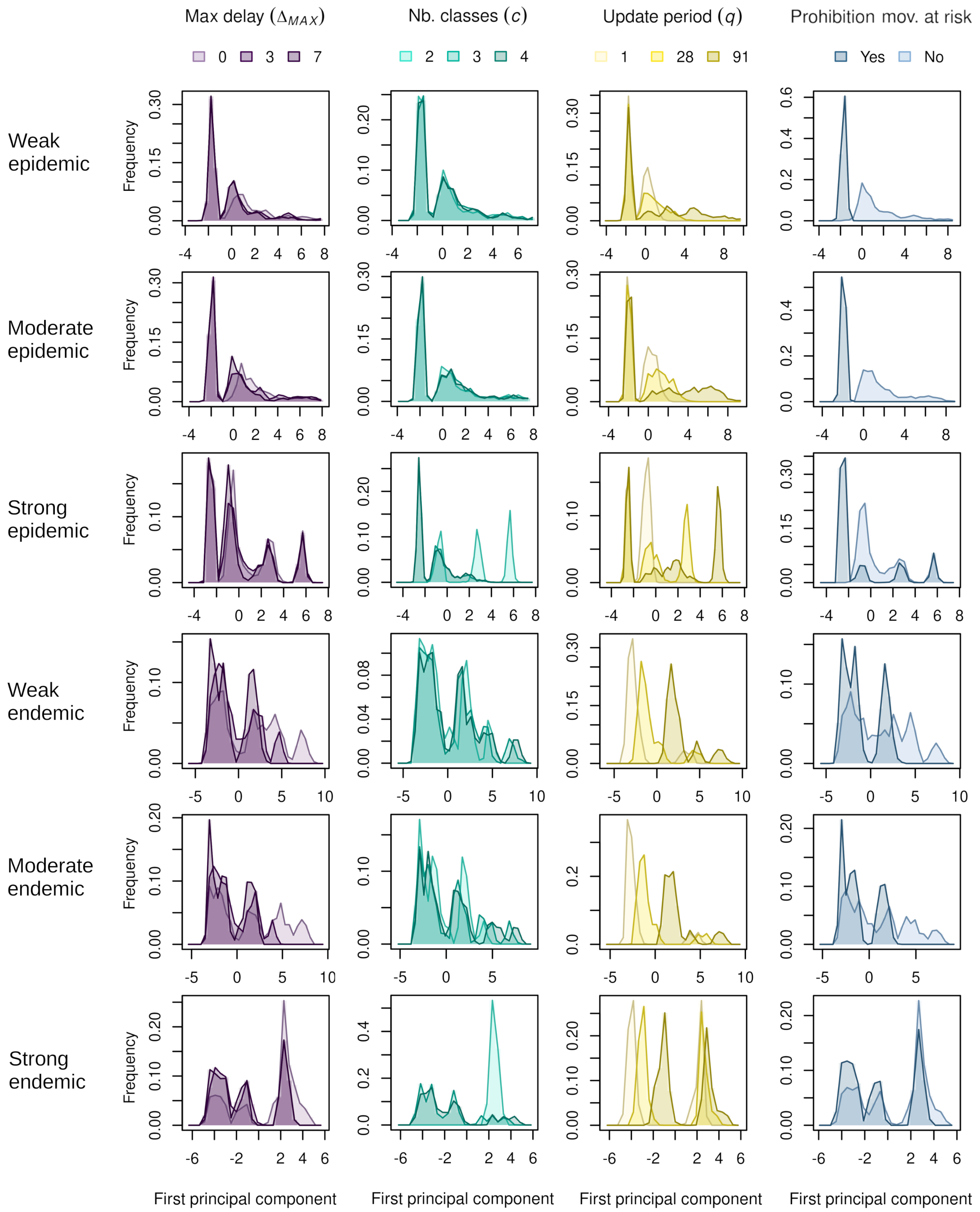


**Figure S6**: Distribution of the simulations on the first principal component of the PCA performed as a first step of the sensitivity analysis, for the six epidemiological settings (rows), according to their algorithm parameter values (columns). The simulations are divided according to their maximal delay (3 shades of purple), their number of prevalence classes (3 shades of cyan), herd status update period (3 shades of yellow) and prohibition of movements at risk (2 shades of blue).

#### 7. Changes in in- and out-degree distributions in the movement network after rewiring

To assess the impact of the algorithm on the in- and out-degrees of the movement network, the distributions of $ind_h$ and $outd_h$ are recorded and their respective percentiles computed for each simulation. The distribution of percentiles for all simulations with the same epidemiological setting (epidemic or endemic) are compared to those of the original, non-rewired network. By definition, $x\%$ of herds have a degree lower or equal to the $x^{th}$ percentile value. In the original dataset, respectively 70% and 45% of the herds don't buy and sell livestock during the year. These proportions could only be modified by the algorithm through the replacement of internal movements by imports and exports, which would only decrease the degrees of the herds in question. Therefore, the following focuses on the distributions between the $70^{th}$ and $100^{th}$ percentile of $ind_h$ and between the $45^{th}$ and $100^{th}$ percentile of $outd_h$.

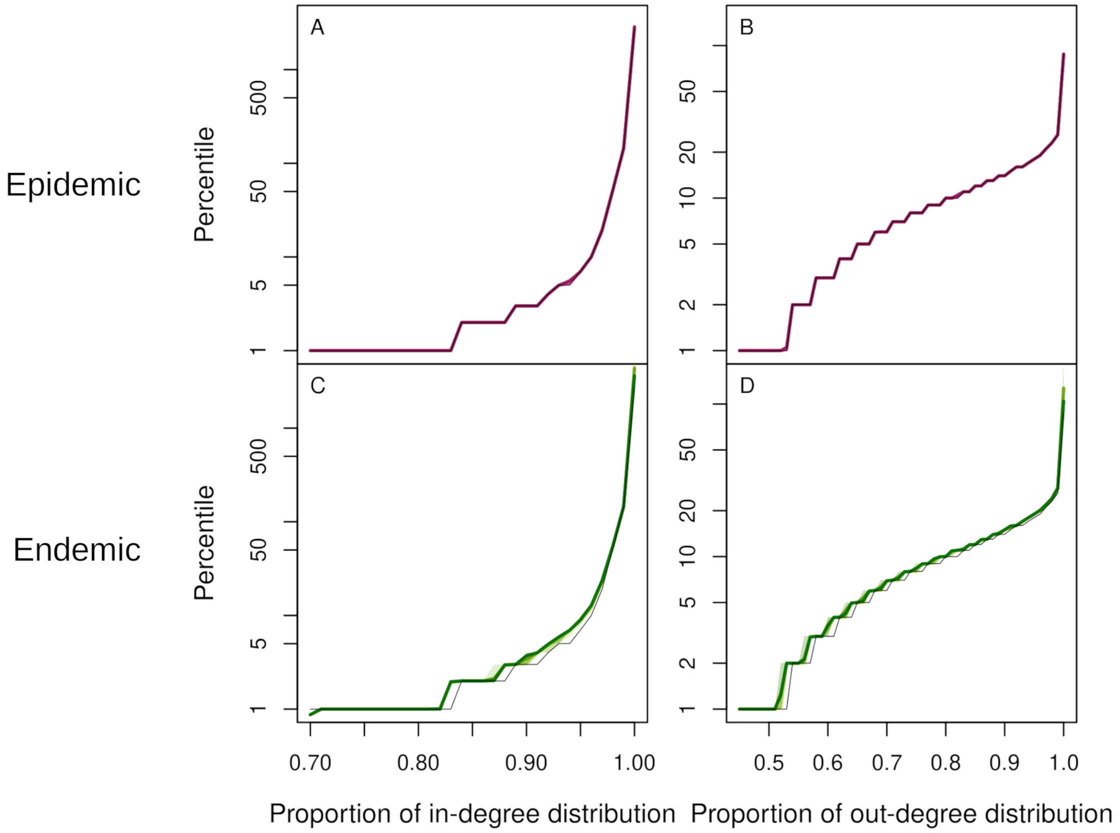


**Figure S7**: Percentiles (in log-scale) of the in-degree $ind_h$ ($1^{st}$ column) and out-degree $outd_h$ ($2^{nd}$ column) distributions after rewiring in epidemic ($1^{st}$ row, magenta) and endemic settings ($2^{nd}$ row, green), compared the distribution those of the original network (black). Null percentiles are not displayed. Each scenario (combination of algorithm parameters) is represented by its average (solid line) and an interval with 80% of simulations (envelope). Given the very low variance between simulations within a same epidemiologic scenario, envelopes are not represented if all 80% of simulations have the same percentile value.

There is no strong difference between the degrees before and after rewiring in epidemic settings. Indeed, Fig. S7A and S7B show that at least 80% of the rewired networks in epidemic settings had degree distributions almost identical to the distribution of the original network, regardless of algorithm parameter values. In endemic settings however, the percentile values are overall higher after rewiring. Indeed, the percentiles of $ind_h$ distribution above the $83^{th}$ are on average 16% higher in rewired network than in the original one (Fig. S7). Similarly, the percentiles of $outd_h$ distribution above the $52^{th}$ are on average 9% higher (S7D). These results indicate a slight increase of the in- and out-degrees of the herds because of rewiring. This increase is evenly distributed across all herd degree levels, indicating that rewiring affects all herds regardless of degree. Besides, Fig. S7C and S7D show that the increase is also observed across all simulations: the envelopes including 80% of the simulations being steadily over the percentile values for the original network indicate that at least 90% of the simulations experience the overall increase in in- and out-degree described above. Even though the increase remains small, it is therefore a consistent impact of the algorithm in endemic settings.
